## Supplementary Information for "Shared pathway-specific network mechanisms of dopamine and deep brain stimulation for the treatment of Parkinson’s disease"

[1]Movement Disorder and Neuromodulation Unit, Department of Neurology, Charité – Universitätsmedizin Berlin, corporate member of Freie Universität Berlin and Humboldt-Universität zu Berlin, Berlin, Germany
[2]Einstein Center for Neurosciences Berlin, Charité – Universitätsmedizin Berlin, corporate member of Freie Universität Berlin and Humboldt-Universität zu Berlin, Berlin, Germany
[3]Bernstein Center for Computational Neuroscience Berlin, Berlin, Germany
[4]Research Group Neural Interactions and Dynamics, Department of Neurology, Max Planck Institute for Human Cognitive and Brain Sciences, Leipzig, Germany
[5]Neurophysics Group, Department of Neurology, Charité – Universitätsmedizin Berlin, corporate member of Freie Universität Berlin and Humboldt-Universität zu Berlin, Berlin, Germany
[6]Berlin Center for Advanced Neuroimaging, Bernstein Center for Computational Neuroscience, Berlin, Germany
[7]Center for Brain Circuit Therapeutics, Department of Neurology, Brigham and Women's Hospital, Harvard Medical School, Boston, MA, USA
[8]Department of Neurosurgery, Massachusetts General Hospital, Harvard Medical School, Boston, MA, USA
[9]Department of Neurosurgery, Charité – Universitätsmedizin Berlin, corporate member of Freie Universität Berlin and Humboldt-Universität zu Berlin, Berlin, Germany
[10]Technische Universität Berlin, Berlin, Germany
[11]Physikalisch-Technische Bundesanstalt Braunschweig und Berlin, Berlin, Germany
[12]NeuroCure Clinical Research Centre, Charité – Universitätsmedizin Berlin, corporate member of Freie Universität Berlin and Humboldt-Universität zu Berlin, Berlin, Germany
[13]Berlin School of Mind and Brain, Humboldt-Universität zu Berlin, Berlin, Germany

### Supplementary Figures

#### Local power in cortex and STN

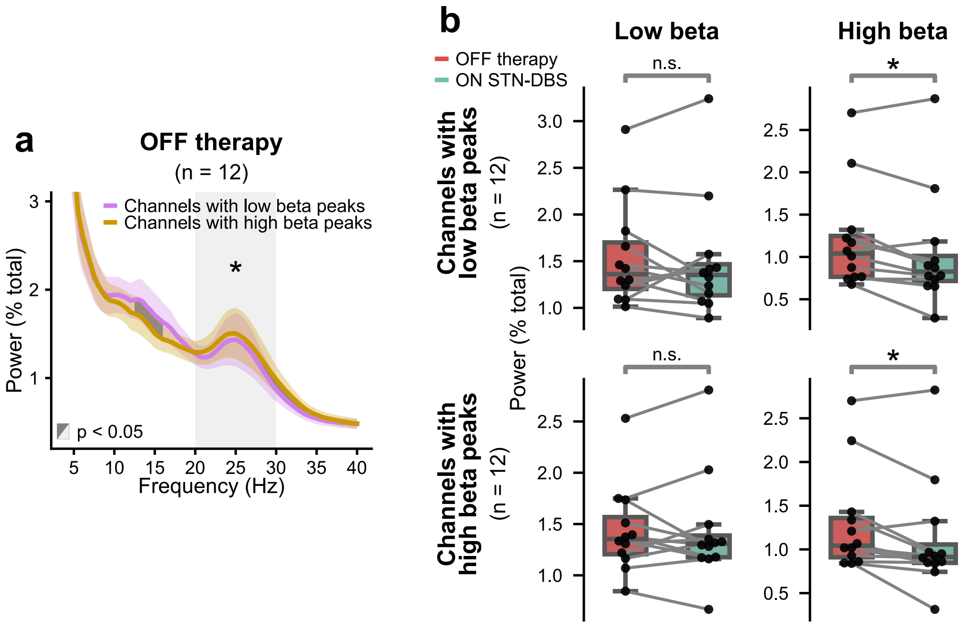


**Supplementary Figure 1: Selection of STN-LFP channels for peaks in low and high beta band power and the effects of DBS.** STN-LFP channels were grouped according to the presence of peaks in power in the low and high beta bands, based on visual inspection of the OFF therapy state. Many channels shared peaks in both the low and high beta bands, and as such were assigned to both groups, leading to some degree of low and high beta activity being present in the opposing groups. **a** The presence of peaks in power was corroborated by a statistical analysis, which showed that the low beta peak group had significantly greater power in the low beta band (12.5-16 Hz; $p < 0.05$, cluster-corrected), while the high beta peak group had significantly greater power in the canonical high beta band ($p < 0.05$). **b** With DBS, no significant reduction in low beta power was found, even in those channels with low beta peaks ($p > 0.05$), whereas significant reductions in high beta power were found in those channels with predominantly low and high beta peaks ($p < 0.05$). * $p < 0.05$. Abbreviations: DBS – deep brain stimulation; n.s. – not significant; STN – subthalamic nucleus.

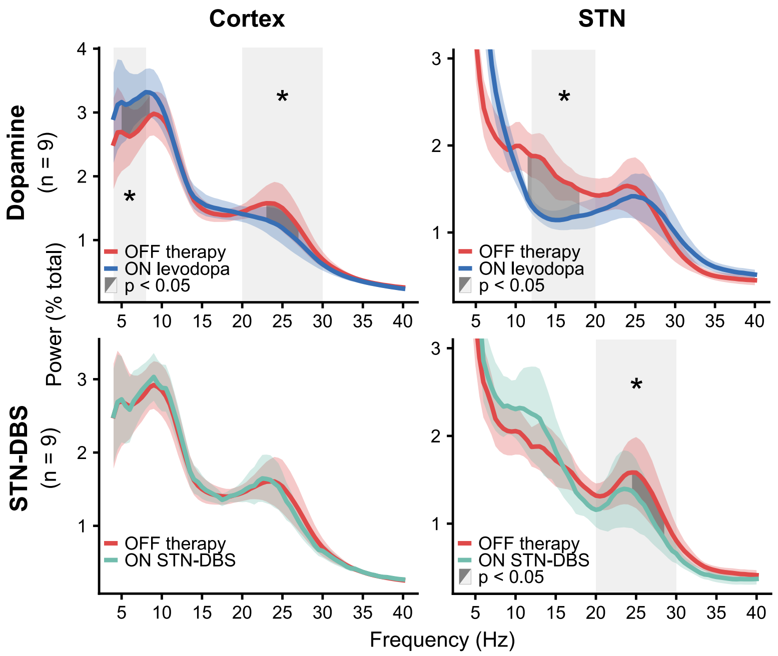


**Supplementary Figure 2: Cortical and subthalamic spectral power in those patients with both OFF therapy – ON levodopa and – ON STN-DBS recordings.** There are distinct modulations in the grand average power spectra across all bipolar contacts in the cortex and STN OFF therapy, ON levodopa, and ON STN-DBS. These modulations strongly resemble those in the wider groups (Fig. 2). Shaded coloured areas show standard error of the mean. Shaded light grey areas indicate a significant difference in the average values of canonical frequency bands between conditions. Shaded dark grey areas indicate clusters of significant differences between conditions for the respective frequency bins. * $p < 0.05$. Abbreviations: DBS – deep brain stimulation; STN – subthalamic nucleus.

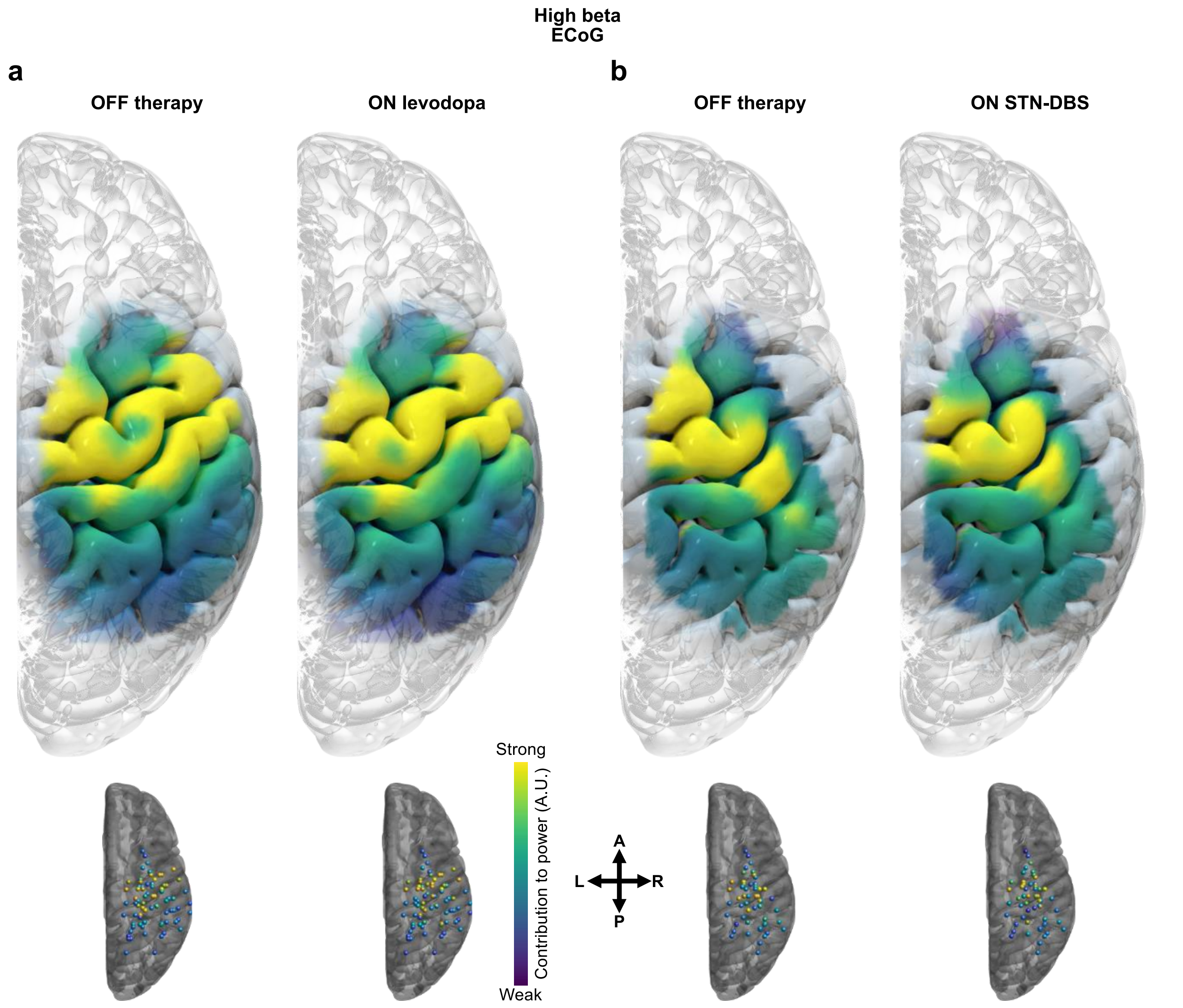


**Supplementary Figure 3: Spatial maps of cortical high beta power.** Localisation of the strongest power component extracted with spatio-spectral decomposition for cortical high beta. Localisations are highly similar across therapeutic states for the **a** OFF therapy – ON levodopa group and **b** OFF therapy – ON STN-DBS group. Localisations are shown for individual electrodes and interpolated to surfaces. Abbreviations: A – anterior; DBS – deep brain stimulation: ECoG – electrocorticography; L – left; P – posterior; R – right: STN – subthalamic nucleus.

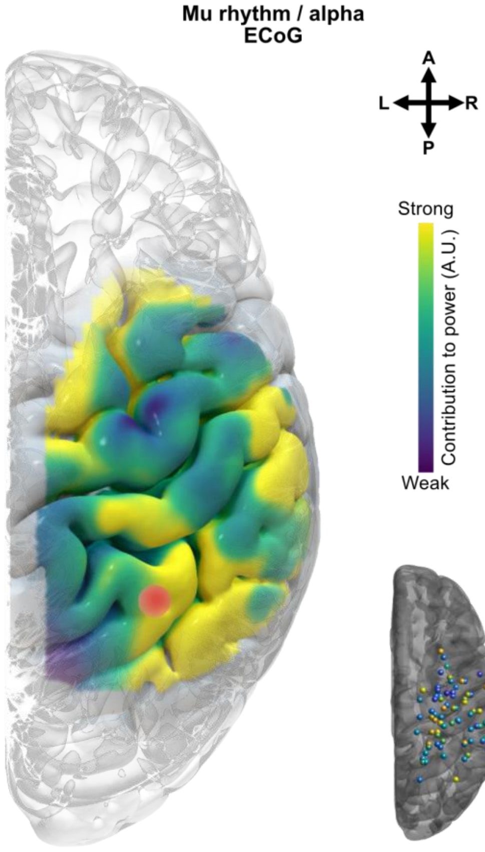


**Supplementary Figure 4: Spatial map of cortical mu rhythm/alpha power.** Localisation of the strongest power component extracted with spatio-spectral decomposition for cortical mu rhythm/alpha, averaged over OFF therapy and ON levodopa. Localisations are shown for individual electrodes and interpolated to surfaces. Red dot shows point of strongest source. Abbreviations: A – anterior; ECoG – electrocorticography; L – left; P – posterior; R – right.

#### Directed sensory cortex – STN communication

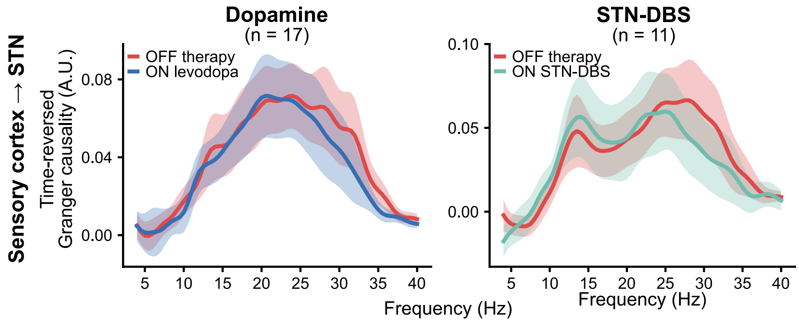


**Supplementary Figure 5: Sensory cortex – STN directed communication.** Granger causality shows sensory cortex drives communication with STN across medication and stimulation states. Shaded coloured areas show standard error of the mean. Note that of the n = 18 (medication) and n = 12 (stimulation) subjects, one subject had no electrode coverage of the sensory cortex, producing the n = 17 and n = 11 group sizes seen here, respectively. Abbreviations: DBS – deep brain stimulation; STN – subthalamic nucleus.

#### Directionality of cortico-subthalamic communication OFF therapy vs. ON STN-DBS

Time-reversed Granger causality (TRGC) revealed the driving force of communication from motor cortex to STN was reduced in the high beta band with STN-DBS (Fig. 3c; Supplementary Fig. 6g).

Given that TRGC is based on net Granger scores, this reduction with stimulation can reflect: 1) reduced information flow from cortex to STN; 2) increased information flow from STN to cortex; or 3) a combination of 1 and 2. To assess which scenario is responsible for the observed change, we can examine the various Granger scores used to compute TRGC. Following the notation of Winkler et al.[1]: $\boldsymbol{X}$ is defined to be the motor cortex seeds, and $\boldsymbol{Y}$ the STN targets; $F_{X\rightarrow Y}$ and $F_{Y\rightarrow X}$ are the Granger scores from seeds to targets and targets to seeds, respectively; $F^{\text{net}}_{X\rightarrow Y}$ is the difference of Granger scores (i.e., $F_{X\rightarrow Y} - F_{Y\rightarrow X}$); $\sim$ represents time-reversal of signals; and finally, $\widetilde{D}^{\text{net}}_{X\rightarrow Y}$ is TRGC (i.e., $F^{\text{net}}_{X\rightarrow Y} - \tilde{F}^{\text{net}}_{\tilde{X}\rightarrow \tilde{Y}}$).

First, $F_{X\rightarrow Y}$ reveals high beta information flow from cortex to STN to be reduced with stimulation (scenario 1; Supplementary Fig. 6a), and strongly resembles the final TRGC scores (Supplementary Fig. 6g). Second, $F_{Y\rightarrow X}$ shows increased information flow from STN to cortex with stimulation from 30 Hz (scenario 2; Supplementary Fig. 6b). Together, these findings would suggest that the change in cortical driving force with stimulation reflects both reduced information flow from cortex and increased information flow from STN (scenario 3). However, the increased information flow from STN to cortex with stimulation from 30 Hz is also present in the time-reversed signals, as $\tilde{F}_{\tilde{Y}\rightarrow \tilde{X}}$ shows (Supplementary Fig. 6e). Given that the purpose of time-reversal is to capture weak data asymmetries – properties of the data which do not reflect causal interactions between signals[2] – we attribute the increased information flow from STN to cortex with stimulation from 30 Hz as 'noise', potentially a residual stimulation artefact given that it continues to ramp towards the stimulation frequency (130 Hz). In contrast, $\tilde{F}_{\tilde{X}\rightarrow \tilde{Y}}$ shows a lack of difference in the high beta band between stimulation conditions (Supplementary Fig. 6d), indicating that the reduced information flow from cortex to STN with stimulation is not the result of weak data asymmetries. Altogether, we attribute the reduced cortical high beta driving force observed in the final TRGC scores to a reduction of information flow from cortex to STN (scenario 1), and not an increase of information flow from STN to cortex.

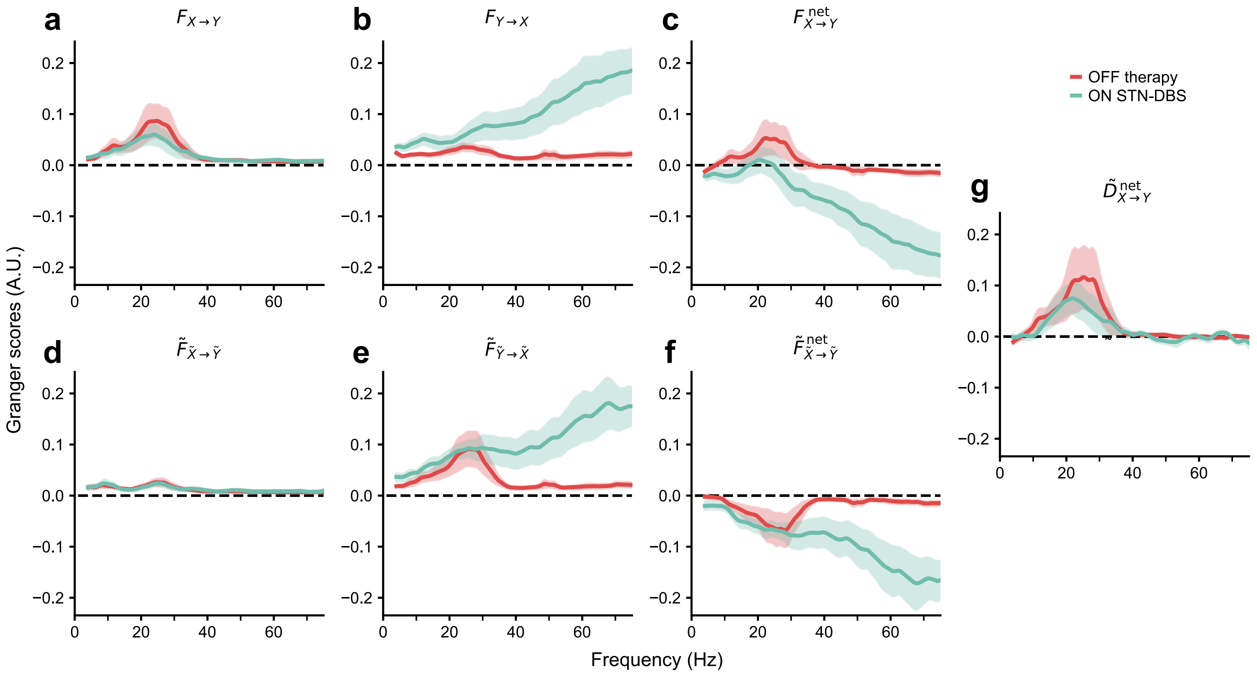


**Supplementary Figure 6: Directionality of cortico-subthalamic connectivity.** We define $\boldsymbol{X}$ to be motor cortex ECoG signals, and $\boldsymbol{Y}$ to be STN-LFP signals. **a** Granger scores from motor cortex to STN. **b** Granger scores from STN to motor cortex. **c** Net Granger scores from motor cortex to STN. **d** Granger scores of time-reversed signals from motor cortex to STN. **e** Granger scores of time-reversed signals from STN to motor cortex. **f** Net Granger scores of time-reversed signals from motor cortex to STN. **g** Time-reversed Granger causality from motor cortex to STN. Shaded coloured areas show standard error of the mean.

#### Spatial maps of cortico-subthalamic coupling

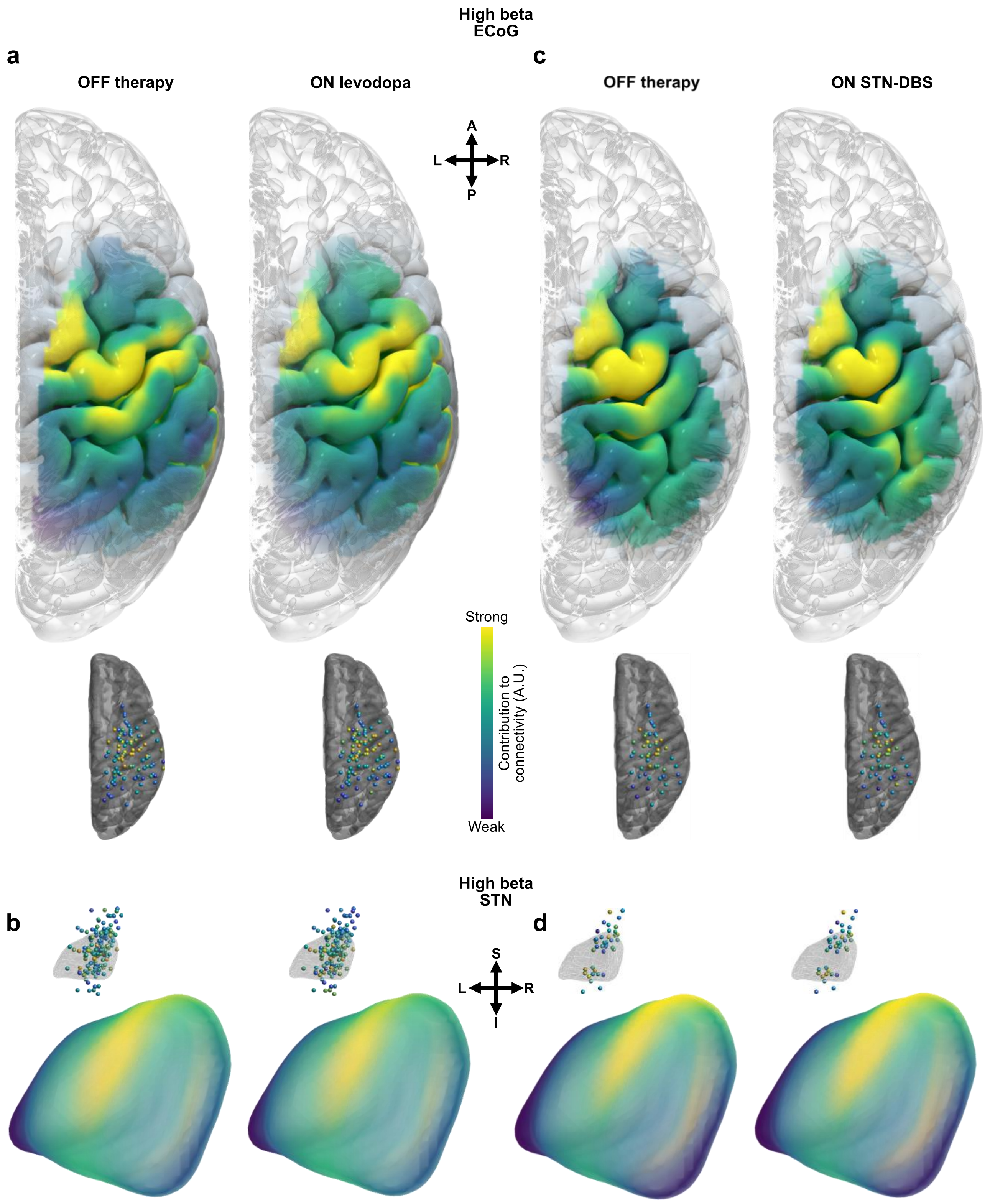


**Supplementary Figure 7: Spatial maps of cortico-subthalamic coupling.** Localisation of the strongest high beta connectivity component extracted from the maximised imaginary coherency analysis. Localisations are highly similar across therapeutic states for the OFF therapy – ON levodopa group in the **a** cortex and **b** STN, as well as for the OFF therapy – ON STN-DBS group in the **c** cortex and **d** STN. Localisations are shown for individual electrodes and interpolated to surfaces. Abbreviations: A – anterior; ECoG – electrocorticography; I – inferior; L – left; P – posterior; R – right; S – superior; STN – subthalamic nucleus.

### Supplementary Tables

#### Subject and recording information

**Supplementary Table 1: Subject information.**

| ID | Age (y) | Sex | DD (y) | LEDD (mg) | Preoperative UPRDS-III (OFF, ON levodopa; A.U.) | 3MFU STN-DBS amplitude (left, right; mA) | 12MFU UPRDS-III (OFF, ON STN-DBS; A.U.) | Preoperative dyskinesia ON levodopa |
|---|---|---|---|---|---|---|---|---|
| EL003 | 56 | F | 15 | 600 | 31, 18 | N/A | N/A | Yes |
| EL004 | 45 | F | 2 | 1462 | n.a. | N/A | N/A | Yes |
| EL006 | 57 | M | 6 | 1000 | 31, 11 | n.a. | 57, 24 | No |
| EL007 | 55 | M | 3 | 850 | 36, 9 | n.a. | 48, 35 | Yes |
| EL008 | 66 | M | 8 | 858 | 59, 28 | N/A | N/A | Yes |
| EL009 | 59 | M | 7 | 1600 | 24, 8 | n.a. | 35, 11 | No |
| EL010 | 66 | F | 12 | 900 | 27, 11 | N/A | N/A | Yes |
| EL011 | 67 | M | 7 | 949 | 30, 9 | N/A | N/A | Yes |
| EL012 | 54 | M | 12 | 866 | 39, 14 | 1.0, 1.7 | 43, 18 | Yes |
| EL013 | 64 | M | 9 | 800 | 31, 12 | N/A | N/A | No |
| EL014 | 52 | M | 14 | 1750 | 35, 8 | 1.5, n.a. | 52, 32 | Yes |
| EL016 | 73 | M | 20 | 1630 | 47, 19 | N/A | N/A | Yes |
| EL017 | 65 | F | 8 | 1150 | 41, 14 | 2.1, 2.3 | n.a. | No |
| EL019 | 58 | M | 10 | 1215 | 39, 18 | 1.7, 2.0 | n.a. | Yes |
| EL020 | 51 | M | 7 | 880 | 32, 24 | N/A | N/A | Yes |
| EL021 | 43 | M | 10 | 1000 | 42, 23 | N/A | N/A | Yes |
| EL022 | 70 | M | 7 | 1073 | 41, 23 | 1.6, 2.0 | n.a. | Yes |
| EL023 | 67 | M | 8 | 1700 | 44, 20 | 1.9, 1.7 | n.a. | Yes |
| EL025 | 65 | M | 14 | 1990 | 58, 25 | 2.8, 2.8 | n.a. | Yes |
| EL026 | 63 | M | 13 | 1225 | 32, n.a. | n.a. | n.a. | Yes |
| EL027 | 32 | M | 5 | 533 | 25, 16 | 1.9, 2.0 | n.a. | Yes |

Note that preoperative UPDRS-III scores were unavailable for subject EL004 and EL026. Note that 3MFU STN-DBS amplitudes were unavailable for subjects EL006, EL007, EL009, EL014 (partially), and EL026. Note that 12MFU UPDRS-III scores were unavailable for subjects EL017, EL019, EL022, EL023, and EL025-27. Average 3MFU STN-DBS amplitudes across hemispheres were: 1.9±0.2 mA (mean±SEM). Abbreviations: DBS – deep brain stimulation; DD – disease duration; F – female; LEDD – levodopa-equivalent daily dose; M – male; N/A – not applicable; n.a. – not available; STN – subthalamic nucleus; UPDRS – Universal Parkinson's Disease Rating Scale; 3MFU – 3-month follow up post-implantation; 12MFU – 12-month follow up post-implantation.

**Supplementary Table 2: Recording information.**

| ID | DBS lead model | ECoG model | ECoG hemi-sphere | Days since implantation (OFF therapy, ON levodopa, ON STN-DBS) | Recording duration (OFF therapy, ON levodopa, ON STN-DBS; s) | Peri-operative STN-DBS amplitude (left, right; mA) |
|---|---|---|---|---|---|---|
| EL003 | MT 3389 | AT TS06R-AP10X-0W6 | Left | 2, 4, N/A | 619, 603, N/A | N/A |
| EL004 | BS Vercise Cartesia | AT TS06R-AP10X-0W6 | Left | 2, 2, N/A | 306, 301, N/A | N/A |
| EL006 | MT SenSight Short | AT TS06R-AP10X-0W6 | Right | 3, 5, 3 | 301, 304, 90 | N/A, 2.0 |
| EL007 | MT SenSight Short | AT DS12A-SP10X-000 | Right | 3, 2, 3 | 311, 196, 334 | 2.0, 1.5 |
| EL008 | BS Vercise Cartesia X | AT TS06R-AP10X-0W6 | Left | 3, 6, N/A | 308, 251, N/A | N/A |
| EL009 | MT SenSight Short | AT TS06R-AP10X-0W6 | Left | 4, 5, 4 | 303, 231, 305 | 2.0, 2.0 |
| EL010 | MT SenSight Short | AT TS06R-AP10X-0W6 | Right | 6, 3, N/A | 306, 231, N/A | N/A |
| EL011 | MT SenSight Short | AT TS06R-AP10X-0W6 | Right | 7, 6, N/A | 306, 316, N/A | N/A |
| EL012 | MT SenSight Short | AT TS06R-AP10X-0W6 | Right | 6, 2, 6 | 307, 306, 307 | 1.5, 1.5 |
| EL013 | MT SenSight Short | AT DS12A-SP10X-000 | Right | 3, 4, N/A | 306, 306, N/A | N/A |
| EL014 | MT SenSight Short | AT TS06R-AP10X-0W6 | Right | 4, 2, 4 | 307, 328, 306 | 2.0, 2.5 |
| EL016 | MT SenSight Short | AT TS06R-AP10X-0W6 | Right | 4, 6, N/A | 306, 306, N/A | N/A |
| EL017 | MT SenSight Short | AT TS06R-AP10X-0W6 | Right | 3, 3, 6 | 237, 241, 306 | 3.0, 2.0 |
| EL019 | MT SenSight Short | AT TS06R-AP10X-0W6 | Right | 5, 5, 6 | 245, 728, 96 | N/A, 2.5 |
| EL020 | MT SenSight Short | AT TS06R-AP10X-0W6 | Right | 3, 2, N/A | 300, 306, N/A | N/A |
| EL021 | MT SenSight Short | AT TS06R-AP10X-0W6 | Right | 2, 3, N/A | 306, 306, N/A | N/A |
| EL022 | MT SenSight Short | AT TS06R-AP10X-0W6 | Right | 4, 2, 4 | 306, 630, 306 | 3.0, 3.0 |
| EL023 | MT SenSight Short | AT TS06R-AP10X-0W6 | Right | 5, 4, 5 | 306, 306, 310 | 2.5, 2.5 |

| | | | | | | |
|---|---|---|---|---|---|---|
| EL025 | MT SenSight Short | AT TS06R-AP10X-0W6 | Right | 6, N/A, 6 | 306, N/A, 306 | 3.0, 3.0 |
| EL026 | MT SenSight Short | AT TS06R-AP10X-0W6 | Right | 3, N/A, 3 | 306, N/A, 306 | 3.0, 2.5 |
| EL027 | MT SenSight Short | AT TS06R-AP10X-0W6 | Left | 3, N/A, 3 | 303, N/A, 306 | 3.0, 3.0 |

Average day of recording post-implantation for each condition was: OFF therapy 3.9±0.3 days; ON levodopa 3.7±0.4 days; ON STN-DBS 4.4±0.4 days (mean±SEM). Average recording duration for each condition was: OFF therapy 314±16 s; ON levodopa 351±37 s; and ON STN-DBS 273±24 s. Average peri-operative STN-DBS amplitudes across hemispheres were: 2.4±0.1 mA. Abbreviations: AT – Ad-Tech; BS – Boston Scientific; DBS – deep brain stimulation; ECoG – electrocorticography; MT – Medtronic; N/A – not applicable; n.a. – not available; STN – subthalamic nucleus.

#### Association of hyperdirect pathway connectivity and oscillatory connectivity

**Supplementary Table 3: Hyperdirect pathway fibres ~ high beta oscillatory connectivity.**

| | Beta coefficient (A.U.) | Standard error (A.U.) | z | p > \|z\| | [0.025 | 0.975] |
|---|---|---|---|---|---|---|
| Intercept | 7.917 | 1.345 | 5.888 | < 0.001 | 5.281 | 10.552 |
| Spatial patterns | 4.064 | 0.850 | -0.088 | < 0.001 | 2.398 | 5.731 |
| C(Medication)[ON] | -0.061 | 0.692 | 4.780 | 0.930 | -1.416 | 1.295 |
| Subject Var | 28.113 | 0.758 | | | | |

$n_{observations}$ = 1676; $n_{groups}$ = 18; mean group size = 93.1 (range 48-168); scale = 200.306; Log-likelihood = -6839.031; Bayesian information criterion = 13707.759; $R^2_{marginal/conditional}$ = 0.037/0.155. Medication state was used as a fixed condition. Subjects were treated as a random effect.

**Supplementary Table 4: Hyperdirect pathway fibres ~ low beta oscillatory connectivity.**

| | Beta coefficient (A.U.) | Standard error (A.U.) | z | p > \|z\| | [0.025 | 0.975] |
|---|---|---|---|---|---|---|
| Intercept | 7.997 | 1.197 | 6.682 | < 0.001 | 5.651 | 10.343 |
| Spatial patterns | -1.142 | 0.790 | -1.446 | 0.148 | -2.689 | 0.406 |
| C(Medication)[ON] | 0.018 | 0.697 | 0.026 | 0.979 | -1.348 | 1.384 |
| Subject Var | 21.298 | 0.574 | | | | |

$n_{observations}$ = 1676; $n_{groups}$ = 18; mean group size = 93.1 (range 48-168); scale = 203.449; Log-likelihood = -6849.839; Bayesian information criterion = 13729.374; $R^2_{marginal/conditional}$ = 0.003/0.097. Medication state was used as a fixed condition. Subjects were treated as a random effect.

#### Association of indirect pathway connectivity and oscillatory connectivity

**Supplementary Table 5: cortex – STN fMRI connectivity ~ low beta oscillatory connectivity.**

| | Beta coefficient (A.U.) | Standard error (A.U.) | z | p > \|z\| | [0.025 | 0.975] |
|---|---|---|---|---|---|---|
| Intercept | -0.048 | 0.003 | -18.297 | < 0.001 | -0.053 | -0.043 |
| Spatial patterns | 0.006 | 0.002 | 2.967 | 0.007 | 0.002 | 0.011 |
| C(Medication)[ON] | < -0.001 | 0.003 | < -0.001 | 1.000 | -0.006 | 0.006 |
| Subject Var | < 0.001 | 0.002 | | | | |

$n_{observations}$ = 214; $n_{groups}$ = 18; mean group size = 11.9 (range 10-12); scale = $5e^{-4}$; Log-likelihood = 489.321; Bayesian information criterion = -957.179; $R^2_{marginal/conditional}$ = 0.063/0.131. Medication state was used as a fixed condition. Subjects were treated as a random effect.

**Supplementary Table 6: cortex – STN fMRI connectivity ~ high beta oscillatory connectivity.**

| | Beta coefficient (A.U.) | Standard error (A.U.) | z | p > \|z\| | [0.025 | 0.975] |
|---|---|---|---|---|---|---|
| Intercept | -0.048 | 0.003 | -18.840 | < 0.001 | -0.053 | -0.043 |
| Spatial patterns | 0.004 | 0.002 | 2.006 | 0.045 | < 0.001 | 0.008 |
| C(Medication)[ON] | < -0.001 | 0.003 | < -0.001 | 1.000 | -0.006 | 0.006 |
| Subject Var | < 0.001 | 0.001 | | | | |

$n_{observations}$ = 214; $n_{groups}$ = 18; mean group size = 11.9 (range 10-12); scale = 5e$^{-4}$; Log-likelihood = 486.911; Bayesian information criterion = -952.359; $R^2_{marginal/conditional}$ = 0.028/0.080. Medication state was used as a fixed condition. Subjects were treated as a random effect.

**Supplementary Table 7: cortex – Putamen, GPe, and STN fMRI connectivity ~ low beta oscillatory connectivity.**

| | Beta coefficient (A.U.) | Standard error (A.U.) | z | p > \|z\| | [0.025 | 0.975] |
|---|---|---|---|---|---|---|
| Intercept | -0.050 | 0.003 | -15.851 | < 0.001 | -0.056 | -0.043 |
| Spatial patterns | 0.005 | 0.002 | 2.271 | 0.023 | 0.001 | 0.010 |
| C(Medication)[ON] | < 0.001 | 0.003 | < 0.001 | 1.000 | -0.006 | 0.006 |
| Subject Var | < 0.001 | 0.003 | | | | |

$n_{observations}$ = 214; $n_{groups}$ = 18; mean group size = 11.9 (range 10-12); scale = 5e$^{-4}$; Log-likelihood = 494.516; Bayesian information criterion = -967.567; $R^2_{marginal/conditional}$ = 0.049/0.219. Medication state was used as a fixed condition. Subjects were treated as a random effect.

**Supplementary Table 8: cortex – Putamen, GPe, and STN fMRI connectivity ~ high beta oscillatory connectivity.**

| | Beta coefficient (A.U.) | Standard error (A.U.) | z | p > \|z\| | [0.025 | 0.975] |
|---|---|---|---|---|---|---|
| Intercept | -0.050 | 0.003 | -16.486 | < 0.001 | -0.056 | -0.044 |
| Spatial patterns | 0.004 | 0.002 | 1.968 | 0.049 | < 0.001 | 0.008 |
| C(Medication)[ON] | < -0.001 | 0.003 | < -0.001 | 1.000 | -0.006 | 0.006 |
| Subject Var | < 0.001 | 0.002 | | | | |

$n_{observations}$ = 214; $n_{groups}$ = 18; mean group size = 11.9 (range 10-12); scale = 5e$^{-4}$; Log-likelihood = 493.632; Bayesian information criterion = -965.799; $R^2_{marginal/conditional}$ = 0.029/0.180. Medication state was used as a fixed condition. Subjects were treated as a random effect.

**Supplementary Table 9: cortex – Putamen and GPe fMRI connectivity ~ low beta oscillatory connectivity.**

| | Beta coefficient (A.U.) | Standard error (A.U.) | z | p > \|z\| | [0.025 | 0.975] |
|---|---|---|---|---|---|---|
| Intercept | -0.050 | 0.003 | -14.986 | < 0.001 | -0.056 | -0.043 |
| Spatial patterns | 0.005 | 0.002 | 2.055 | 0.040 | < 0.001 | 0.010 |
| C(Medication)[ON] | < -0.001 | 0.003 | < -0.001 | 1.000 | -0.006 | 0.006 |
| Subject Var | < 0.001 | 0.003 | | | | |

$n_{observations}$ = 214; $n_{groups}$ = 18; mean group size = 11.9 (range 10-12); scale = 5e$^{-4}$; Log-likelihood = 484.166; Bayesian information criterion = -946.869; $R^2_{marginal/conditional}$ = 0.039/0.218. Medication state was used as a fixed condition. Subjects were treated as a random effect.

**Supplementary Table 10: cortex – Putamen and GPe fMRI connectivity ~ high beta oscillatory connectivity.**

| | Beta coefficient (A.U.) | Standard error (A.U.) | z | p > \|z\| | [0.025 | 0.975] |
|---|---|---|---|---|---|---|
| Intercept | -0.050 | 0.003 | -15.484 | < 0.001 | -0.056 | -0.044 |
| Spatial patterns | 0.004 | 0.002 | 1.842 | 0.065 | < -0.001 | 0.008 |
| C(Medication)[ON] | < -0.001 | 0.003 | < -0.001 | 1.000 | -0.006 | 0.006 |
| Subject Var | < 0.001 | 0.002 | | | | |

$n_{observations}$ = 214; $n_{groups}$ = 18; mean group size = 11.9 (range 10-12); scale = $5e^{-4}$; Log-likelihood = 483.579; Bayesian information criterion = -945.693; $R^2_{marginal/conditional}$ = 0.025/0.188. Medication state was used as a fixed condition. Subjects were treated as a random effect.
